# Single-cell transcriptional analysis reveals naïve helper ILC-like cells in zebrafish

**DOI:** 10.1101/342477

**Authors:** Pedro P. Hernández, Paulina M. Strzelecka, Emmanouil I. Athanasiadis, Ana F. Robalo, Catherine M. Collins, Pierre Boudinot, Jean-Pierre Levraud, Ana Cvejic

## Abstract

Innate lymphoid cells (ILCs) are important mediators of the immune response and homeostasis in barrier tissues of mammals. However, the existence and function of ILCs in other vertebrates is poorly understood. Here, we use single-cell RNA sequencing to generate a comprehensive atlas of zebrafish lymphocytes during tissue homeostasis and following immune challenge. We profiled 14,080 individual cells from the gut of wild-type zebrafish, as well as of *rag1*-deficient fish which lack T and B cells, and discovered diverse populations of helper ILC-like cells. Unexpectedly, fish displayed a *rorc*-positive, naïve subset that established a Type 3 or Type 2 response only upon immune challenge. Specifically, naïve ILC-like cells expressed *il22* and *tnfa* following exposure to inactivated bacteria, or *il13* following exposure to helminth extract. Cytokine-producing ILC-like cells express a specific repertoire of novel immune-type receptors, likely involved in recognition of environmental cues. We identified additional novel markers of zebrafish ILCs and generated a cloud repository for their in-depth exploration.

## Introduction

Vertebrate immune systems consist of the innate arm, which responds immediately to challenge, and the adaptive arm, which responds via acquired antigen receptors. In mammals, myeloid cells (granulocytes, mast cells, monocytes/macrophages, dendritic cells) form the innate immune system, whereas B and T lymphocytes contribute to the adaptive immune response (Boehm, 2012; Riera Romo et al., 2016). Recently discovered innate lymphoid cells (ILCs) represent a rare population of lymphocytes (Artis and Spits, 2015; Eberl et al., 2015; Walker et al., 2013). Unlike T and B cells, ILCs do not express antigen receptors or undergo clonal expansion when stimulated. Instead, in the absence of adaptive antigen receptors, ILCs sense environmental cues mostly through cytokine receptors, and promptly respond to signals by producing distinct cytokines. During homeostasis, humans and mice contain four populations of ILCs: natural killer (NK) cells, and three subsets of helper ILCs (ILC1, ILC2 and ILC3). NK cells bear similarity to cytotoxic T cells (CD8+ cell), which directly kill cells infected with intracellular pathogens. Helper ILCs in human and mouse are classified as ILC1, ILC2 and ILC3 based on their transcription factor (TF) and cytokine secretion profiles, as well as phenotypic cell-surface markers (Artis and Spits, 2015; Eberl et al., 2015; Spits et al., 2013; Walker et al., 2013). Both Th1 and ILC1 express T-bet (encoded by *tbx21*) as well as so-called “Th1 cytokines” such as interferon gamma (IFNγ) and tumour necrosis factor alpha (TNFα), and act against intracellular pathogens. Th2 and ILC2 express GATA-3 and secrete IL-4, IL-13 and amphiregulin, and contribute to defence against helminths and venoms. Th17 and ILC3s express RORγt (encoded by *RORC*) as well as IL-17a, IL-17f and IL-22 and promote immunity against extracellular bacteria and fungi (Annunziato et al., 2015; Artis and Spits, 2015; Eberl et al., 2015; Simoni et al., 2017). Apart from NK cells, ILCs have been characterized only in mouse and human (Artis and Spits, 2015; Eberl et al., 2015).

The different immune cell types are usually distinguished based on expression of specific CD (cluster of differentiation) markers. However, a homogenous population of blood cells, as defined by surface markers, may include many distinct transcriptional states with different functional properties (Guo et al., 2013; Jaitin et al., 2014; Villani et al., 2017; Wilson et al., 2015). In addition, the surface markers used to define distinct human and murine leukocyte subsets are not the same, making it difficult to compare cell types across different species. Therefore, there is a need for unbiased methodologies that define immune cell types based on cellular state rather than cell surface markers. This is particularly relevant for species other than mouse and human, where specific antibodies for distinct blood and immune cell types are not readily available.

In zebrafish, the heterogeneity of haematopoietic cells has mostly been investigated with fluorescent transgenic reporter lines, as very few antibodies for surface markers are available (Carradice and Lieschke, 2008). These approaches have confirmed the presence of erythrocytes, thrombocytes, neutrophils, macrophages, eosinophils, T cells, B cells and NK cells in zebrafish. Comprehensive transcriptome atlases exist for many of these cell types (Athanasiadis et al., 2017; Carmona et al., 2017; Tang et al., 2017). Because these studies focused on steady state conditions, they were limited in their ability to characterise the response mechanisms following immune challenge. Compared to mice and humans, little is known about the diversity of T cells and cytokine-producing innate lymphoid cells in zebrafish, and a detailed characterization of their transcriptional profiles is still lacking. As a result, the evolution of T cells and ILCs in non-mammalian species remains ambiguous.

Here, we characterized the repertoire of innate and adaptive lymphocytes in zebrafish. Using single-cell RNA sequencing, we generated a comprehensive atlas of cellular states of lymphocytes collected from various organs in steady state and following immune challenge. We discovered cytokine-expressing ILC subsets, including a “naïve” population of ILCs that converted to ILC2- or ILC3-like cells in response to specific immune challenge. Thus, unlike human and mice which already have diverse helper ILC subsets during homeostasis, fish contain naïve helper ILCs that diversify only upon immune challenge. Further, in contrast to mammalian ILC subsets which respond to environmental signals via cytokine receptors, our data suggest that the “naïve” ILCs in zebrafish sense environmental cues through novel immune-type receptors, akin to natural cytotoxicity receptors. Finally, we generated a cloud repository for the community to access our sequencing data.

## Results

### Single cell transcriptional analysis of lymphocytes in zebrafish

Lymphocyte development occurs through several stages, at distinct body locations, each marked with specific gene expression patterns. To capture the diversity of lymphoid cell types, we purified and sequenced the RNA from single cells collected from primary lymphoid organs (kidney and thymus), secondary lymphoid organs (spleen) as well as barrier tissues (gut and gills) of healthy, unstimulated adult zebrafish. We used three different transgenic lines: *Tg(lck:EGFP*) (Langenau et al., 2004), which labels T cells as well as NK cells (Carmona et al., 2017); *Tg(cd4-1:mCherry)* (Dee et al., 2016), which labels CD4 T cells and macrophages, and *Tg(mhc2dab:GFP, cd45:dsRed)* (Wittamer et al., 2011) (Figure 1A) that is expected to label B cells (when sorted as GFP+/DsRed-).

**Figure 1.**
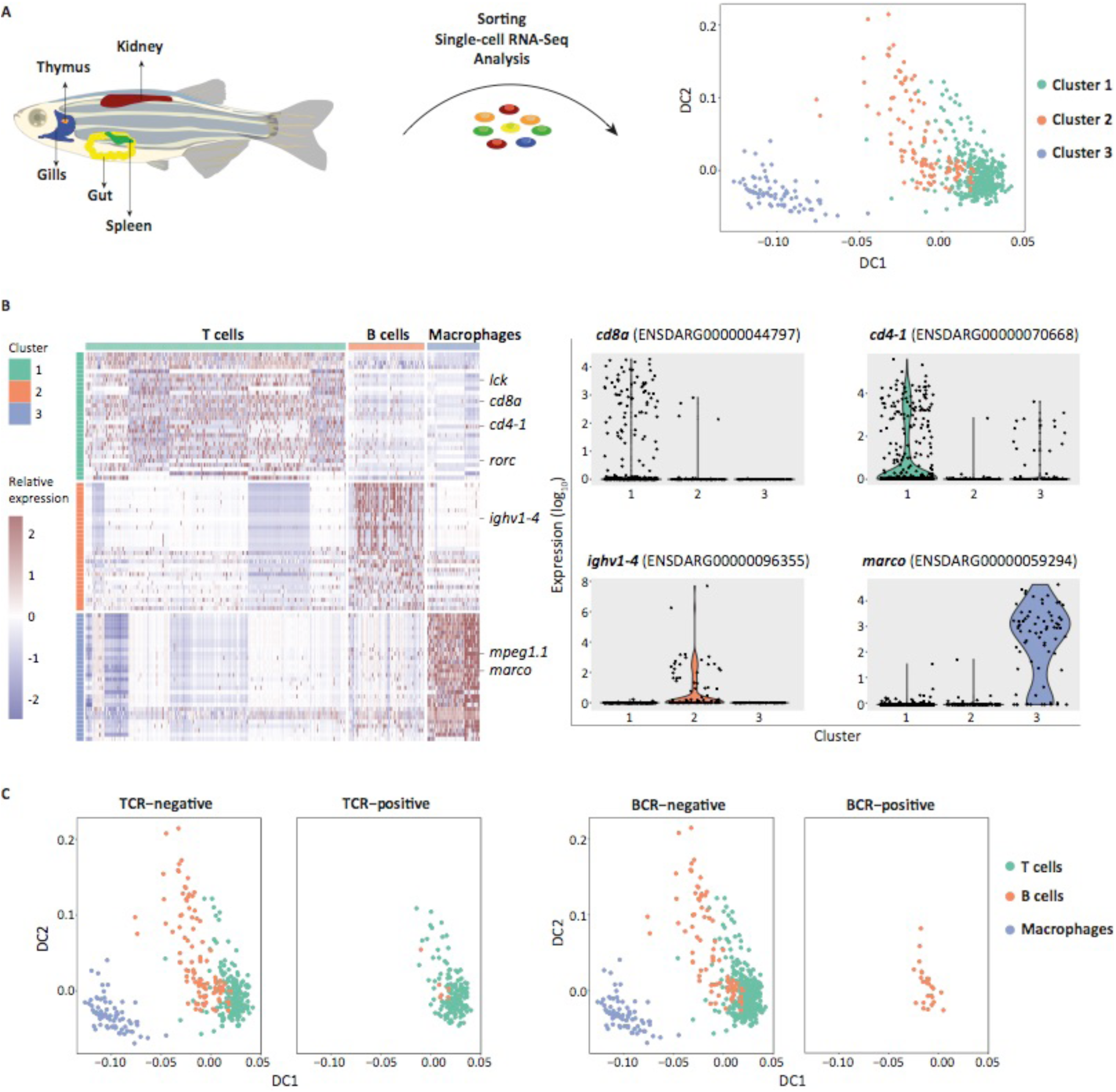
Identification of immune cell types in zebrafish during steady state haematopoiesis. **A.** Experimental strategy for sorting single cells from transgenic zebrafish lines. Cells were collected from different organs and transgenic lines, namely, Tg(cd4-1:mCherry), Tg(lck:EGFP) and Tg(mch2dab:GFP, cd45:DsRed). Single cells were index-sorted into 96-well plates and processed with standard Smart-seq2 protocol followed by computational analysis. **B.** Heatmap showing the expression level of marker genes for each of the identified cluster. Columns represent cells and rows marker genes. Violin plots showing the expression of signature genes in each cluster. **C.** TraCeR and BraCeR analysis showing cells positive and negative for TCR and BCR rearrangements across the cell types.

We first performed single-cell RNA sequencing (Smart-seq2) of reporter-positive cells; 542 cells out of 796 passed quality control (QC) and were subjected to further analysis (Supp. Figure 1). Based on 3,374 highly variable genes (HVGs) inferred from biological cell-to-cell variation (Supp. Figure 2A), we generated Diffusion Maps and clustered cells within the 3D diffusion space. Our hierarchical clustering approach revealed three main populations.

The cells in the first cluster (C1) appeared to be T cells with high expression of *cd4-1*, *cd8a* and *lck* (Figure 1B). As expected, cells in this cluster originated from *cd4-1:mCherry* and *lck:EGFP* transgenic cells collected from kidney, gills, gut, thymus and spleen (Supp. Figure 2B). T cell identity is defined by the expression of a T cell receptor (TCR) and the successful rearrangement of TCR genes is essential for the cell survival during T-cell development in the thymus. To further confirm our computational prediction that cells in C1 are indeed T cells, we applied TraCeR (Stubbington et al., 2016), a novel method for reconstruction of TCR sequences from single-cell RNA-seq data. We were able to unambiguously detect V(D)J recombination events in 224 cells (Figure 1C, Supp. Table 1). Incidence of V(D)J recombination in TCR was highly associated with cluster C1, which provided additional evidence of their T cells identity (Figure 1C, Supp. Table 1). Importantly, we did not observe T cell clonal expansion (i.e. multiple cell with the same rearrangements), which is expected to occur when the fish are exposed to pathogens.

The second cluster (C2) had a signature of B lymphocytes and cells in this cluster originated from kidneys of the *Tg(mhc2dab:GFP, cd45:dsRed)* line (Supp. Figure 2B). They showed expression of immunoglobulin-heavy variable 1-4 (*ighv1-4*), an orthologue of human immunoglobulin heavy constant mu gene (*IGHM*) (Figure 1B). Importantly, we detected BCR rearrangements in 36 cells from this cluster using BraCer, therefore confirming their B cell identity (Lindeman et al., 2017) (Figure 1C, Supp. Table 1), again with no detectable clonal expansion.

The cluster three (C3) was exclusively comprised of cells that originated from *cd4-1:mCherry* transgenic cells collected from gills, gut and spleen (Supp. Figure 2B). These cells had a high expression of macrophage receptor with collagenous structure (*marco*) and macrophage expressed gene 1 (*mpeg1.1*), (Figure 1B) strongly indicative of their macrophage identity. This was not surprising, as *cd4-1:mCherry* has been found to label both CD4 T cells and macrophages (Dee et al., 2016).

Thus, single-cell RNA-Seq of *lck:EGFP*+, *mhc2dab:GFP+/cd45:dsRed-* and *cd4-1:mCherry+* cells identified the adaptive lymphocytes in zebrafish, namely T and B cells, and their transcriptional signatures.

### Identification of immature thymocytes, Tregs and CD8 cells in zebrafish

Cells in C1 were collected from thymus, kidney, gills, gut, and spleen of *lck:EGFP*+ and *cd4-1:mCherry*+ lines and thus provided a unique opportunity to assess differentiation of T lymphocytes in zebrafish (Figure 2). We generated diffusion maps to place all T cells into pseudo-time and reconstructed their development in an unbiased way (Figure 2). We observed a differentiation trajectory from double-positive (DP) thymocytes, that showed high expression of *rag1*, *rorc*, *cd4-1* and *cd8a* to single-positive (SP) thymocytes that downregulated *rag1* and *rorc* but displayed exclusive expression of *cd4-1* or *cd8a*. These data strongly suggest that thymocyte maturation is highly conserved between mammals and zebrafish.

**Figure 2.**
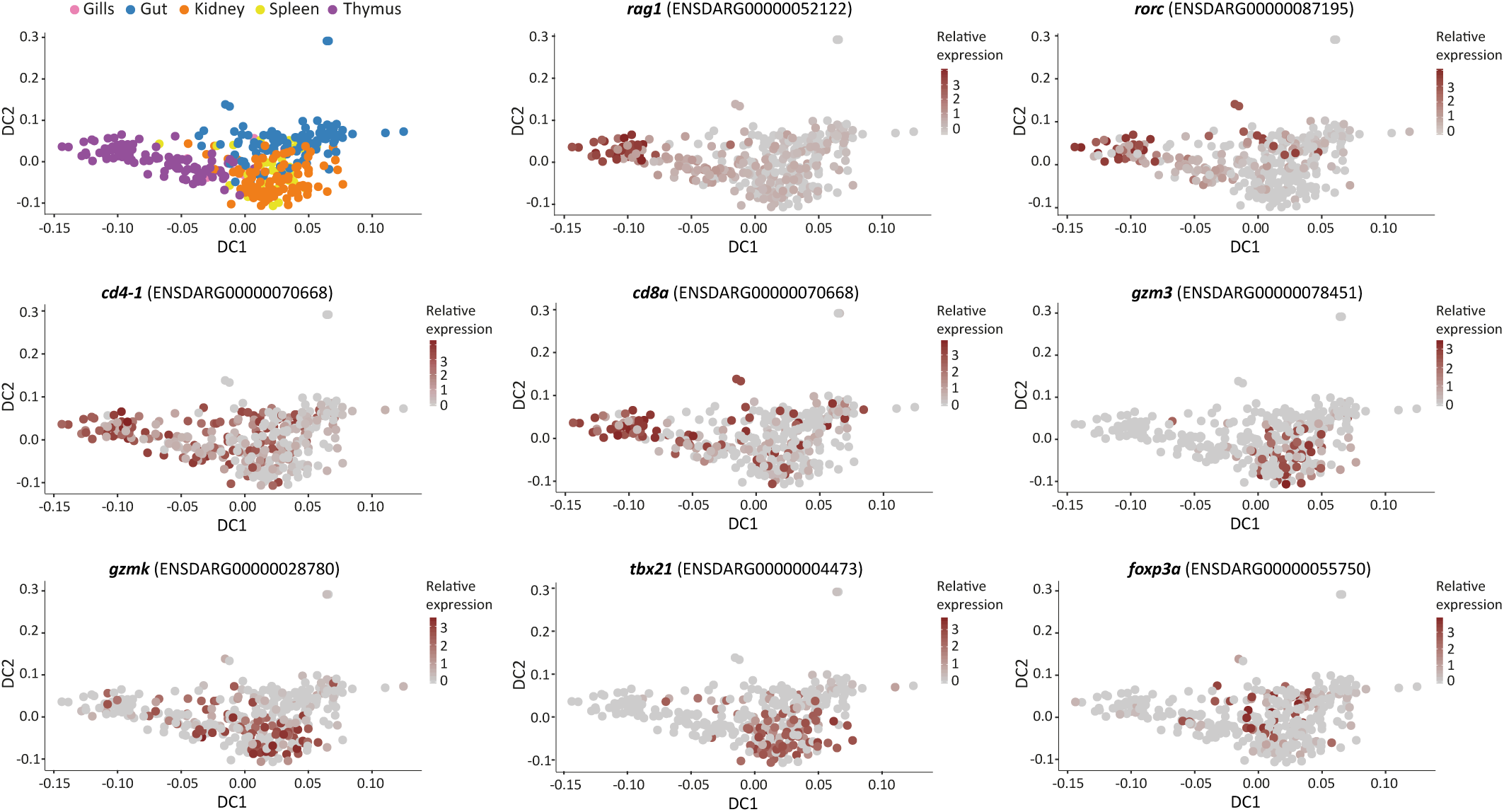
Differentiation trajectory of T cells in zebrafish. Diffusion maps of 337 T cells collected from the gills, gut, kidney, spleen and thymus coloured by the tissue of origin (top, left panel) and expression level in log2(counts+1) scale of signature genes (all other panels).

The pseudo-time trajectory also captured a heterogeneous population of mature T cells. Conventional T cells in mammals and fish are subdivided according to the mutually exclusive expression of the TCR co-receptors - CD4 for helper T cells and CD8 for cytotoxic T cells (Wang and Bosselut, 2009). In mammals, cytotoxic CD8+ T cells (CTLs) kill infected/dysfunctional cells by releasing perforins, granzymes, and granulysin (Dotiwala et al., 2016). Indeed, we identified cells that had high expression of *cd8a*, granzymes (*gzm3* and *gzmk*) and *tbx21* (master TF of CTL in mammals shown to promote cytotoxicity of CD8+ cells) (Lazarevic et al., 2013) (Figure 2). Conversely, we identified a population of cells that expressed *cd4-1*, possibly labelling helper T cells. Within the population of CD4+ cells a few cells expressed *foxp3a*, suggesting the presence of regulatory T cells (Tregs) (Kasheta et al., 2017) (Figure 2).

### *rag1^−/−^* mutants lack T and B cells but have cytokine-producing cells in the gut

Rag1- and Rag2-deficient mouse strains, which lack adaptive but retain innate lymphoid cells (Fort et al., 2001; Hurst et al., 2002; Sawa et al., 2011), have provided substantial insight into ILCs. These mice showed expression of many cytokines previously considered to be T cell-specific and therefore provided the first evidence of the existence of helper ILCs (Fort et al., 2001; Hurst et al., 2002; Sawa et al., 2011). Thus, to focus on innate lymphoid cells in zebrafish, we turned to *rag1*^−/−^ fish. As in mice, *rag1*^−/−^ zebrafish lack T and B lymphocytes (Wienholds et al., 2002) but retain NK cells (Petrie-Hanson et al., 2009).

In line with previous reports (Petrie-Hanson et al., 2009; Tokunaga et al., 2017; Wienholds et al., 2002), *rag1^−/−^* fish displayed a reduced population of lymphoid cells in the gut as defined by FSC/SSC gating on FACS (Figure 3A-B). Further, bulk qPCR on FACS-sorted cells from the lymphoid population of *rag1*^−/−^ fish showed two- and four-fold decreases in the expression of T cell markers such as *cd3z* and *trac*, respectively, compared to the wild-type fish, whereas the expression level of the pan-lymphocyte markers *il7r* and *lck* remained the same (Figure 3C). To verify that *rag1*^−/−^ fish indeed lack adaptive lymphocytes, we sequenced 171 *lck:EGFP+* single cells collected from gut and kidney of the *rag1^−/−^* fish and searched for V(D)J recombination events in individual cells. No TCR rearrangements were detected in cells isolated from *rag1^−/−^* fish (Supp. Table 1). These data confirm that the *rag1*^−/−^ provides an excellent tool to examine the innate lymphocytes in zebrafish.

**Figure 3.**
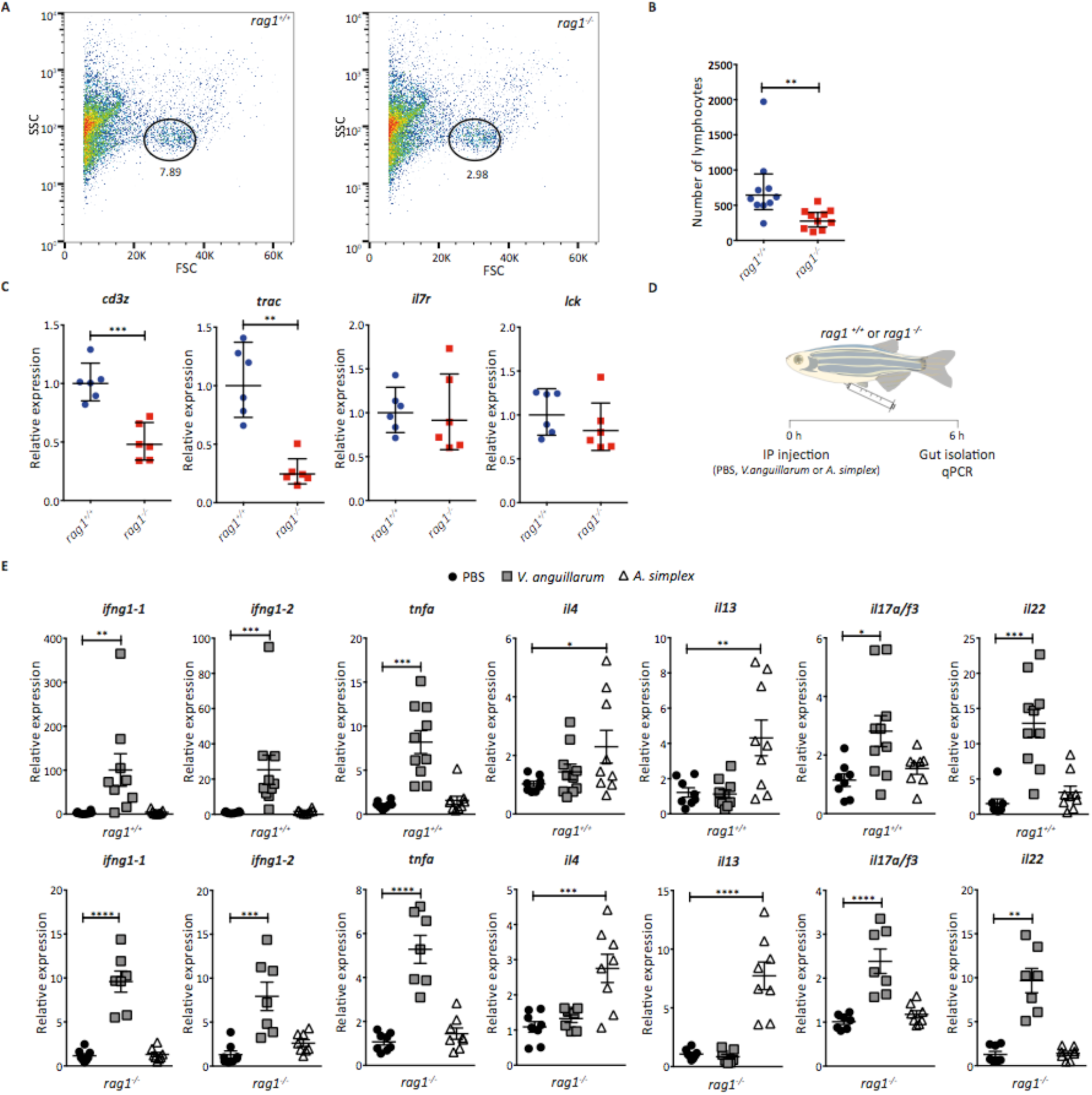
Rag1^−/−^ zebrafish have cytokine producing cells in the gut. **A.** FACS plots showing the percentage of lymphocytes (as defined by FSC/SSC gating) in the gut of wild-type fish (left) and rag1^−/−^ mutant (right). **B.** Number of lymphocytes in the gut of wild-type and rag1^−/−^ mutant fish, 50 000 events recorded. Bars represent the geometric mean ± 95% confidence interval to estimate total number of lymphocytes. Mann-Whitney test. **C.** qPCR expression of T cells associated markers (cd3z, trac) and pan-lymphocyte markers (il7r, lck) in mutant and wild-type fish. Bars represent the geometric mean ± 95% confidence interval to estimate fold changes. Mann-Whitney test. **D.** Scheme of short-term inflammation experiment. **E.** qPCR expression of immune type 1 (ifng1-1, ifng1-2), immune type 2 (il4, il13) and immune type 3 (il17a/f3, il22) signature cytokines in the gut of the wild-type (rag1^+/+^) and mutant (rag1^−/−^) fish following six hours challenge with V. anguillarum or A. simplex. Bars represent the geometric mean ± 95% confidence interval to estimate fold changes. One-way ANOVA test.

Mammals contain three populations of helper ILCs (ILC-1, ILC-2 and ILC-3) that rapidly respond to different tissue signals by producing effector cytokines (Chang et al., 2011; Hernández et al., 2015; Takatori et al., 2009). To study this process in zebrafish, we established short-term inflammation models that trigger cytokine expression of potential ILCs in zebrafish gut (Figure 3D). Formalin-inactivated *Vibrio anguillarum* has been used as a fish vaccine and is known to induce type 3 immunity (Corripio-Miyar et al., 2009); whereas the nematode *Anisakis simplex*, a common fish parasite, is expected to induce type 2 immunity (Nieuwenhuizen et al., 2006). We injected wild-type and *rag1*^−/−^ fish intraperitoneally with PBS (control), or extracts of inactivated *V. anguillarum* or of lyophilised *A. simplex*. Six hours post-injection, we dissected the guts and evaluated the expression of signature cytokines by qRT-PCR (Figure 3D, E). We found that, in both wild-type and *rag1*^−/−^ fish, injection of *V. anguillarum* extract induced the expression of Th1/ILC1 cytokines such as *ifng1-1*, *ifng1-2* as well as Th17/ILC3 cytokines, *il17a/f3* and *il22* (Figure 3E). The expression levels of the Th2/ILC2 cytokines *il4* and *il13* remained unchanged in *V. anguillarum* fish compared to PBS-injected fish. Conversely, injection of *A. simplex* extract induced the expression of Th2/ILC2 cytokines *il4* and *il13* but not of *ifng1-1*, *ifng1-2*, *tnfa*, *il17a/f3* and *il22* (Figure 3E).

These findings have two important implications. First, they confirm that intraperitoneal injection of *V. anguillarum* extract induces a type 1/type 3 immune response in zebrafish gut, and of *A. simplex* extracts induce a type 2 immune response. Second, they reveal the presence of cytokine-producing cells in the gut of immune-challenged *rag1^−/−^* fish, in the context of T cell deficiency. Given that mammalian ILCs have phenotypes that mirror polarized Th subsets in their expression of effector cytokines, our data suggest that the gut in zebrafish contains *bona fide* ILC subtypes.

### Identification of ILC2- and ILC3-like cells in zebrafish

ILCs comprise around 0.5-5% of lymphocytes in barrier tissues in mammals and as such represent a rare population of cells (Halim and Takei, 2014; Simoni et al., 2017). As the *LCK* gene is expressed in all three ILC subtypes in humans (Supp. Figure 3), we reasoned that its expression pattern could be conserved in zebrafish. To capture ILCs subtypes in zebrafish, we utilised our short-term inflammation protocol on *Tg(lck:EGFP) rag1^−/−^* fish. Single-cell RNA sequencing of thousands of *lck:EGFP+* cells isolated from a gut of immune-challenged *rag1^−/−^* mutants provided a powerful approach to study cytokine-producing ILCs in zebrafish.

10x Genomics captures single cells in droplets, such that 5000 cells can be captured and subsequently sequenced within a single run (Zheng et al., 2017). As above, we injected *Tg(lck:EGFP) rag1^−/−^* mutant zebrafish intraperitoneally with PBS, inactivated *V. anguillarum* or lyophilised *A. simplex* extracts and sorted *lck:EGFP+* cells from the gut six hours post-injection. To ensure that a sufficient number of cells were loaded on 10x, we combined an equal number of *lck:EGFP*+ cells for each condition (PBS, *A. simplex* and *V. anguillarum*) from three different fish (nine fish in total). By using this approach, we generated a comprehensive data set that included 3211 single *lck:EGFP+* cells from the guts of PBS-injected, 3626 cells from *A. simplex*-injected and 3487 cells from *V. anguillarum*-injected *rag1^−/−^* zebrafish (Figure 4). On average, we detected 600 genes per cell. Clustering, followed by unbiased identification of marker genes for each cluster (see Methods section), revealed ILC-like cells in all three datasets (Figure 4, Supp. Figure 4, Supp. Figure 5A).

**Figure 4.**
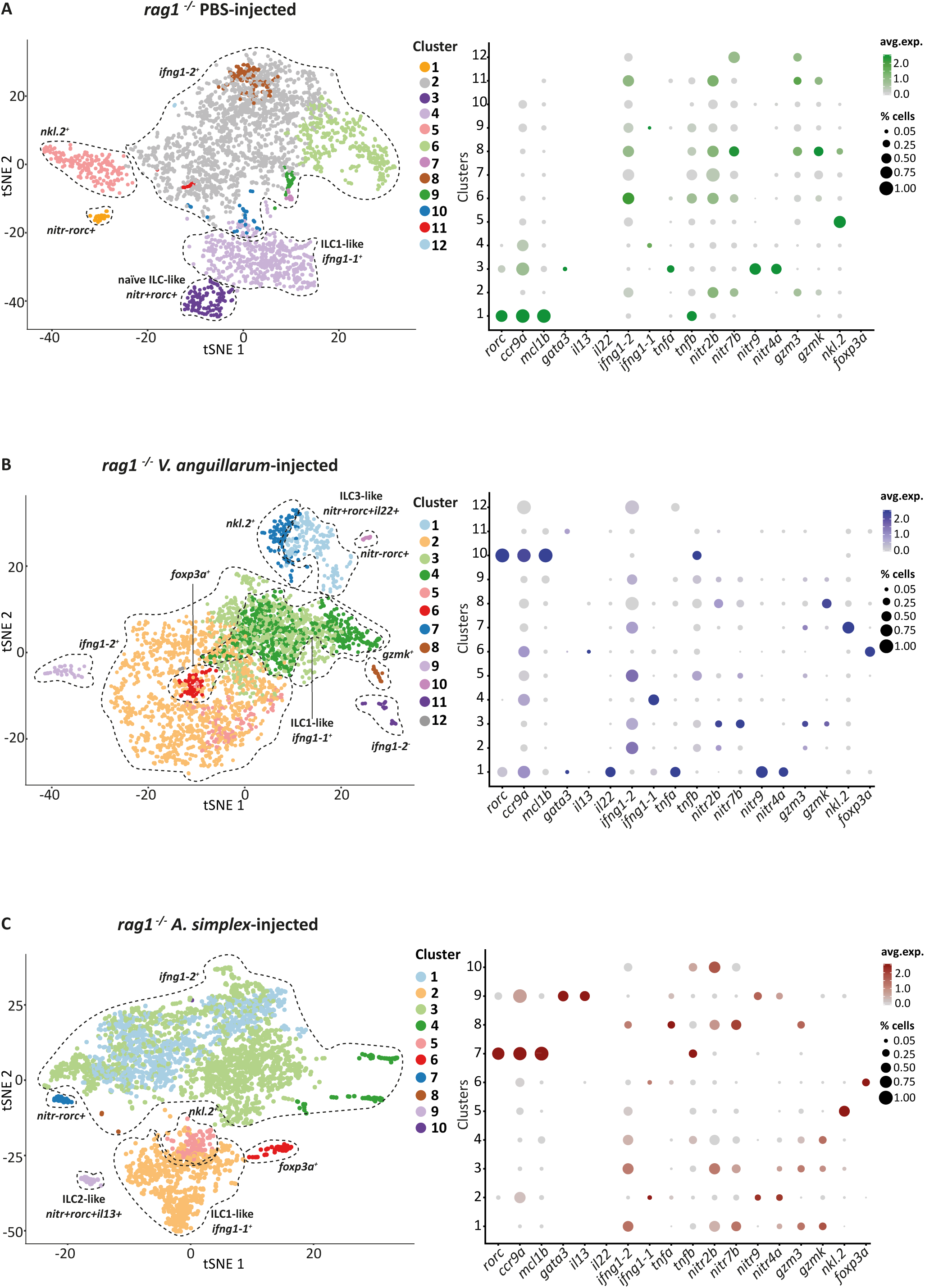
Analysis of the lck+ cells, collected from the gut of rag1^−/−^ fish. 2D projection of tSNE analysis of 10x RNAseq data showing heterogeneity of innate lymphoid cells. Dotplots show the level of expression of marker genes and percentage of cells per cluster that express the gene of interest.

We first analysed *lck:EGFP+* cells from PBS-injected fish and identified several clusters of lymphocytes that expressed *ifng1-2* and granzyme genes (*gzm3*, *gzmk*) (Figure 4A). This transcriptional signature resembles mammalian NK cells. In addition, we identified cells that exclusively expressed *ifng1-1* but not granzymes or nk lysins (cluster 4, Figure 4A). Cells in cluster 4 could be considered ILC1-like cells in zebrafish.

Importantly, our analysis revealed two rare (0.8% and 3.8%) populations of *rorc*+ cells (clusters 1 and 3 on Figure 4A and Supp. Figure 4). The TF RORγt (encoded by *RORC*) is expressed at low levels in circulating and tissue resident ILC precursors in human, as well as in mature ILC3 in human and mouse (Lim et al., 2017a; Scoville et al., 2016). *RORC* is required for the development and function of ILC3 (Sanos et al., 2009; Satoh-Takayama et al., 2008). We found that the *rorc+* clusters did not express cytotoxicity-associated genes such as granzymes (*gzm3*, *gzmk*) or nk lysins (*nkl.2*). To investigate the potential function of these two *rorc+* clusters, we performed differential expression (DE) analysis followed by gene ontology (GO) enrichment analysis (see Methods). We found that *rorc*+ cluster 1 was associated with GO-terms like “response to stress” and “protein folding” (Supp. Figure 5B) and showed high expression of the pro-survival gene *mcl1b*, transcription factor *sox13,* and tumour necrosis factor beta (*tnfb*), an orthologue of human lymphotoxin alpha (*LTA*), but was negative/low for novel immune-type receptor genes (e.g. *nitr2b*, *nitr7b*, *nitr9*, *nitr4a*, etc.), as well as *ifng1-2/ifng1-1* (from here on *nitr-rorc+* cluster) (Figure 4, Supp. Figure 4). In contrast, *rorc*+ cluster 3 was associated with terms like “immune system process”, “response to interferon gamma” and “response to other organisms” (Supp. Figure 5B) and was positive for tumour necrosis factor alpha (*tnfa*) as well as novel immune-type receptors *nitr9* and *nitr4a*, but also negative for *ifng1-2/ifng1-1* (from here on *nitr+rorc+* cluster) (Wei et al., 2007; Yoder et al., 2010) (Figure 4A, Supp. Figure 4). Cells within this cluster also expressed *gata3*.

In *V. anguillarum*-injected fish, *nitr+rorc+* cells (cluster 1), but not *nitr-rorc+* cells (cluster 10), expressed *il22* and elevated *tnfa* (Figure 4B). In humans and mice, ILC3 cells produce the cytokines IL-22 and TNFα cells upon stimulation with bacteria, triggering antimicrobial response and repair programs in epithelial cells during infection (Lindemans et al., 2015; Wolk et al., 2004; Zheng et al., 2008). These data strongly suggest the existence of ILC3-like cells in the zebrafish gut that respond to immune challenge by producing relevant cytokines.

Interestingly, in fish injected with lyophilised *A. simplex*, *nitr+rorc+* cells (cluster 9) expressed *il13* and elevated *gata3* (Figure 4C, Supp. Figure 4). The *gata3* TF is highly expressed in ILC2 cells and is required for their development (Hoyler et al., 2012); it plays a critical role in activating IL-13 production in ILC2 upon stimulation, thus promoting anti-helminth immunity. Therefore, these cells (Figure 4C) resemble mammalian ILC2 cells. Again, *il13*-producing cells were *nitr+* but lacked expression of granzymes, *nkl.2* and *ifng1-2*.

Immune challenged fish (*V. anguillarum*- or *A. simplex*-injected) also had a population of *foxp3a*+ cells (Figure 4B-C, Supp. Figure 4). Cells in this cluster were negative for interferon gamma genes, *nitr* genes and granzymes as well as *cd4-1*. It is tempting to speculate that these cells might represent the zebrafish equivalent of recently reported mammalian regulatory innate lymphoid cells (Wang et al., 2017). However, it should be noted that we did not detect expression of *il10* in this cluster.

To validate that the *rorc*+ cells are a genuine constituent of the zebrafish gut, and not only present in *rag1*^−/−^ fish, we sequenced and analysed additional 3756 GFP+ cells from the gut of PBS-injected wild-type *lck:EGFP* fish. As expected, most of these cells were T cells that expressed *cd8a* or *cd4-1* (specifically Tregs). However, we also identified two clusters of *rorc+* cells that were negative for T cell marker genes: namely, *nitr-rorc+* cells that expressed *mcl1b* and *tnfb,* and *nitr+rorc+* cells that expressed *tnfa* (Supp. Figure 6). These cells represented 0.8% and 1.9% of the *lck:EGFP*+ cell population of wild-type fish. Taken together, our transcriptional profiling of innate lymphocytes in *rag1*-deficient and wild type fish identified ILC-like populations.

### Response of zebrafish ILC-like cells to immune challenge

Our dataset revealed distinct, heterogeneous populations of innate lymphoid cells within the guts of PBS, *V. anguillarum*-, and *A. simplex*-injected fish. We hypothesized that these distinct populations are biologically relevant, reflecting the response of immune cells to disparate stimuli. To test this hypothesis, we evaluated whether the distinct cell types identified in control fish matched those present in *V. anguillarum*- and *A. simplex*-injected fish, and whether the specific treatments led to transcriptional changes in these cell types. We performed integrated analysis of PBS-, *V. anguillarum-,* and *A. simplex*-injected fish using a recently developed computational strategy for scRNA-seq alignment (Butler et al., 2018). This methodology was specifically designed to allow comparison of RNA-Seq datasets across different conditions (Figure 5A). By following the Seurat alignment workflow, we uncovered “shared” cell types across all three datasets and compared their gene expression profiles.

**Figure 5.**
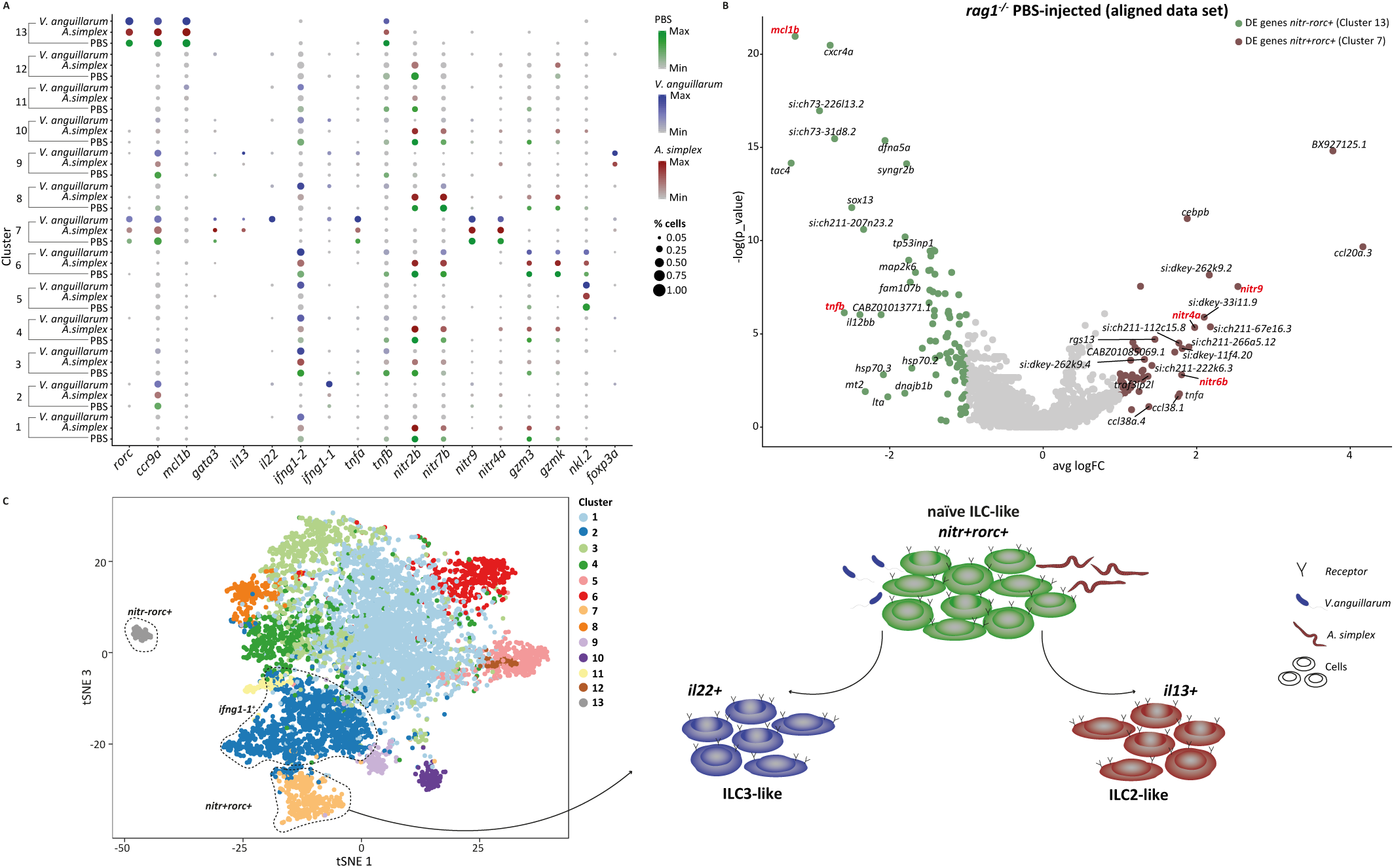
Integrated analysis of PBS - V. anguillarum - A. simplex-injected rag1^−/−^ fish. **A.** Dotplot with the expression level of selected marker genes in each of the clusters. The size of the dots indicates the percentage of cells within the cluster that express the gene of interest; each cluster contains cells from three different conditions **B.** Volcano plot showing the top 20 differentially expressed genes between nitr+rorc+ (Cluster 7) and nitr-rorc+ (Cluster 13) cells originated from rag1^−/−^ PBS-injected fish using aligned dataset **C.** 2D tSNE projection of canonical correlation analysis (CCA) of merged 10x datasets from PBS, V. anguillarum- and A. simplex-injected rag1^−/−^ fish. Model illustrating the strategy that nitr+rorc+ cells use to recognise environmental cues and to commit to ILC2- or ILC3-like cells.

Our analysis showed that the *nitr+rorc+* cells found separately in the three conditions (clusters 3, 1 and 9 on Figures 4A-C, respectively) corresponded to a unique cell subtype (cluster 7 on figure 5A; Supp. Figure 7A). Cells in this cluster upregulated *il22, rorc and tnfa* following *in vivo* stimulation with inactivated *V. anguillarum* (Figure 5A, Supp. Figure 7B). In contrast, injection of *A. simplex* resulted in upregulation of *il13* and *gata3* (Figure 5A), but *ifng1-1, ifng1-2*, *tnfb* and *nitr* genes remained unaltered relative to the control (Figure 5A). These data suggest the presence of a “naïve” population of ILCs (*nitr+rorc+)* in zebrafish that differentiate into either ILC2- or ILC-3 like cells only upon immune challenge, depending on the distinct stimuli they receive (Figure 5C).

Similarly, the *nitr-rorc+* population of cells identified separately in PBS-, *V. anguillarum*- and *A. simplex*-injected fish (clusters 1, 10 and 7 on Figures 4A-C, respectively) grouped as distinct cluster (cluster 13 in Figure 5A). These cells expressed *mcl1b*, *sox13* and *tnfb* (orthologue of human lymphotoxin alpha, *LTA*) (Figure 5A, B). *MCL1* is pro-survival gene relevant for maintenance of viability but not of proliferation and is often expressed in long-lived cells (Sathe et al., 2014; Vikström et al., 2016; Yang-Yen, n.d.); whereas the human *tnfb* ortholog *LTA* is expressed in lymphoid tissue inducer (LTi) cells and is involved in the regulation of cell proliferation, differentiation and survival. Unlike “naïve” ILCs, *nitr-rorc+* cells did not respond to immune stimuli by expressing cytokines (Figure 5B, Supp. Figure 7B).

We next asked whether unstimulated *nitr+rorc+* cells express unique surface receptors that enable them to respond to the immune challenge. In addition to *nitr9* and *nitr4a,* “naïve” ILCs specifically expressed novel immune-type receptors *nitr6b* and *nitr5* as shown by DE analysis (Figure 5B). In contrast with human unstimulated ILC subsets, less than 10% of unstimulated *nitr+rorc+* cells expressed cytokine receptors, toll-like and other pattern recognition receptors, suggesting that ILC-like cells in zebrafish recognize environmental cues through natural cytotoxicity receptors (Figure 5C). The developmental origins and hierarchical relationship between *nitr-rorc+* and *nitr*+*rorc*+ populations, however, remains unclear.

Finally, cells identified in cluster 4 in PBS-injected fish, cluster 4 in *V. anguillarum*-injected fish and cluster 2 in *A. simplex*-injected fish (Figures 4A-C, respectively) grouped as cluster 2 (Figure 5A). These cells showed clear upregulation of *ifng1-1* following immune challenge with *V. anguillarum* and no expression of granzymes or nk lysins. These data further support that these cells potentially represent ILC1-like cells in zebrafish.

Altogether, our analyses of over 10,000 single cells collected from the gut of *rag1^−/−^* fish identified previously unappreciated diversity of innate lymphoid cells in zebrafish and revealed how this heterogeneity translates to cell-specific immune responses.

### Cloud repository

To allow easy retrieval of sequencing data from zebrafish innate and adaptive lymphocytes we generated a cloud repository with transcriptional profiles of over 14,000 single cells collected from healthy and immune challenged zebrafish. Together with the data set we previously published on myeloid cells, this is the most comprehensive transcriptional atlas of blood cell types in zebrafish to date (https://www.sanger.ac.uk/science/tools/lymphocytes/lymphocytes/).

### Discussion

Our work provides a comprehensive atlas of both adaptive and innate lymphocytes across different organs in healthy and immune challenged zebrafish. The differentiation trajectory of T cells, showing transition from DP to SP thymocytes and their maturation to heterogeneous populations including CD8+ T cells and Tregs in the kidney, gut, gills and spleen, suggest that T cell maturation is strongly conserved. Importantly, we identified populations of innate lymphocytes in *rag1*-deficient and wild-type fish that resemble helper ILC subtypes in mammals. By analysing 14,080 *lck:EGFP*+ single cells collected from gut of unstimulated and stimulated fish, we discovered two previously unknown populations of *rorc+* ILC-like cells in zebrafish, *nitr+rorc+* and *nitr-rorc+*, which appear in some ways to recapitulate NCR+ ILC3 and NCR- ILC3 subsets, respectively, in humans and mice.

We obtained functional insight into these two distinct populations of *rorc*+ cells by exposing adult zebrafish to specific stimuli that rapidly induce corresponding cytokines in the gut. The population of “naïve” ILC-like cells (*nitr+rorc+*) can convert into ILC3-like cells via upregulation of *il22, tnfa* and *rorc* following stimulation with inactivated *V. anguillarum*, or into ILC2-like cells via upregulation of *il13* and *gata3* following exposure to *A. simplex*. Although this differs somewhat from conventional classification of ILCs, ILCs in mammals can co-express cytokines of more than one type and are capable of transiting from one subtype to another by changing the expression of relevant TFs and cytokines (Gronke et al., 2016; Lim et al., 2017b). Therefore, this plasticity in expression of cytokines and TFs in ILCs appears to be a feature acquired early on in their evolution.

Circulating and tissue resident ILC precursors (ILCPs) in human and mouse also express RORC at low level, and differentiate into multiple ILC subtypes *in vitro* (Lim et al., 2017a; Scoville et al., 2016). However, unlike naïve ILCs in zebrafish, ILCPs cannot be stimulated to produce effector cytokines; only already differentiated ILC1, ILC2 and ILC3 respond to immune challenge by producing relevant cytokines (Lim et al., 2017a; Scoville et al., 2016). This is an important distinction between ILCPs identified in mouse and human compared to the naïve ILCs that we found in zebrafish: naïve ILCs in zebrafish respond to immune challenge promptly, within six hours, and likely transiently, and for these reasons should not be considered a progenitor population.

The survey of cell surface receptors also suggested significant differences in the way ILC-like cells in zebrafish respond to external stimuli compared to their mammalian counterparts. Whereas both human and mouse ILCs constitutively express receptors for cytokines and are mainly activated by cytokines released by the epithelium or antigen presenting cells, zebrafish naïve ILC-like cells express cytokine receptors in less than 10% of cells. More recently, other receptors such as aryl hydrocarbon receptor, Toll-like receptors, and other pattern recognition receptors, as well as natural cytotoxicity receptors (NCRs) have been reported to enable mouse and human ILC3 cells to directly sense environmental cues and induce cytokine expression (Glatzer et al., 2013; Killig et al., 2014). Again, zebrafish ILC-like cells did not express the orthologues of these receptors with the exception of novel immune type receptors (NITRs). Teleost genomes contain multiple NITR genes which are considered to be the functional homologues of mammalian NCRs and KIRs (Yoder et al., 2010). Since naïve ILCs expressed a specific subset of *nitr* genes, we hypothesize that cytokine induction in these cells is triggered by recognition of environmental cues via these receptors.

In addition to ILC3 cells, another population of RORC-expressing ILCs has been identified in mammals - Lymphoid tissue inducer cells (LTis) (Mebius et al., 1997). LTis contribute to the formation of lymph nodes and gut-associated lymphoid tissue, including Peyer’s patches, isolated lymphoid follicles and cryptopatches. Zebrafish gut does not have such organized lymphoid structures (Brugman, 2016) and consequently it is presumed that they don’t have LTis (Lane et al., 2012). In our datasets, we identified a population of *nitr-rorc+* cells that exclusively expressed *tnfb* (a zebrafish orthologue of human lymphotoxin (*LTA*), a marker gene for LTi cells). Although *nitr-rorc+* share some transcriptional features with LTis it is hard to speculate on the function *of nitr-rorc+* cells in zebrafish and whether they are evolutionarily linked with mammalian LTi cells. Interestingly, zebrafish *nitr-rorc+* cells expressed *mcl1b* which is pro-survival gene relevant for maintenance of viability but not of proliferation and is often expressed in long-lived cells (Sathe et al., 2014; Vikström et al., 2016; Yang-Yen, n.d.).

We also identified *ifng* producing cells. Zebrafish have two ifng genes (*ifng1-1* and *ifng1-2*) which show mutually exclusive expression in our data set. The products of two *ifng* genes bind to different receptors in zebrafish and are thus functionally specialized (Aggad et al., 2009). The *ifng1-2* was co-expressed with granzymes or granulysins in the population transcriptionally resembling NK-like cells. In contrast, *ifng1-1* expressing cells had no expression of granzymes nor granulysins and thus represent a distinct subtype of innate lymphocytes, possibly ILC1-like cells. Similar to human ILCs from tonsils, *ifng* producing cells showed no expression of the Th1 master regulator T-bet.

Further studies are necessary to resolve developmental origins, functional properties and potential hierarchical relationship between zebrafish ILC-like subsets. Like humans and mice, zebrafish appear to contain an established ILC1-like population that responds to immune challenge. However, in contrast to mammals, zebrafish display population of naïve ILCs that have ability to convert to *il22* or *il13* producing ILCs only following immune challenge. This discovery could illuminate ILC biology and shed light on their diversification in mammals.

## Methods

### Zebrafish strains and maintenance

The maintenance of zebrafish wild-type line (AB), transgenic lines *Tg(lck:EGFP)* (Langenau et al., 2004), *Tg(cd4-1:mcherry)* (Dee et al., 2016), *Tg(mhc2dab:GFP, cd45:dsRed)* (Wittamer et al., 2011) and *rag1^hu1999^* mutants (also known as *rag1^t26683/26683^* (Wienholds et al., 2002)) was performed in accordance with EU regulations on laboratory animals. *Hu1999* mutation results in a premature stop codon in the middle of the catalytic domain of the Rag1 protein and is considered a null allele (Wienholds et al., 2002).

### FACS sorting

Kidneys from heterozygote transgenic fish either wild-type or *rag1*^−/−^ mutant, were dissected and processed as previously described (Athanasiadis et al., 2017). The guts, spleens, gills and thymuses were dissected and placed in ice cold PBS/5% foetal bovine serum. Single cell suspensions were generated by first passing through a 40 µm strainer using the plunger of a 1 ml syringe as a pestle. These were then passed through a 20 µm strainer before adding 4’,6-diamidino-2-phenylindole (DAPI, Beckman Coulter, cat no B30437) to the samples. For Smart-seq2 experiment individual cells were index sorted into 96 well plates using a BD Influx Index Sorter. Cells from kidney, gut, gills, spleen and thymus from non-transgenic zebrafish line were used for gating.

For the 10x experiment, guts from either *Tg(lck:EGFP) rag1*^−/−^ mutant or wild-type fish were isolated and single cell suspensions were prepared as described above. Three fish, per each condition (i.e. fish intraperitoneally injected with PBS, lyophilised *Anisakis simplex* or inactivated *Vibrio anguillarum*), were used to collect the total of 12,000 lck+ cells (4000 per fish) for 10x experiment.

### Plate-based single-cell RNA processing

The Smart-seq2 protocol (Picelli et al., 2013) was used for whole transcriptome amplification and library preparation as previously described. Generated libraries were sequenced in pair-end mode on Hi-Seq4000 platform.

### Droplet-based single-cell RNA processing

Following the sorting, cells were spun down and resuspended in ice cold PBS with 0.04% bovine serum albumin at the concentration of 500 cells/μl. Libraries were constructed using Chromium™ Controller and Chromium ™ Single Cell 3’ Library & Gel Bead Kit v2 (10x Genomics) according to the manufacturer’s protocol for 5000 cells recovery. Briefly, cellular suspension was added to the master mix containing nuclease-free water, RT Reagent Mix, RT Primer, Additive A and RT Enzyme Mix. Master mix with cells was transferred to the wells in the row labelled 1 on the Chromium ™ Single Cell A Chip (10x Genomics). Single Cell 3’ Gel Beads were transferred into the row labelled 2 and Partitioning Oil was transferred into the row labelled 3. The chip was loaded on Chromium ™ Controller to generate single-cell GEMs. GEM-RT was performed in a C1000 Touch Thermal cycler (Bio-Rad) at the following conditions: 53°C for 45 min, 85°C for 5 min, held at 4°C. Post GEM-RT cleanup was performed with DynaBeads MyOne Silane Beads (Thermo Fisher Scientific). cDNA was amplified using C1000 Touch Thermal cycler at the following conditions: 98°C for 3 min, 12 cycles of (90°C for 15 s, 67°C for 20 s and 72°C for 1 min), 72°C for 1 min, held 4°C. Amplified cDNA was cleaned with the SPRIselect Reagent Kit (Beckman Coulter) and quality was assessed using 2100 Bioanalyser (Agilent). Libraries were constructed following the manufacturer’s protocol and sequenced in pair-end mode on Hi-Seq4000 platform.

### Short-term inflammation experiments

*Vibrio anguillarum* strain 1669 was grown in TSA broth medium to OD600 1.5. Bacterial pellet (9 mL of full grown culture) was resuspended in NaCl 9 g/L, 0.35% formaldehyde, and incubated overnight at 20°C. The suspension was washed four times in NaCl 9g/l and resuspended in 800 μl of the same isotonic solution.

*Anisakis simplex* larvae, extracted from wild herring (*Clupea harengus*), were lyophilized using a freeze dryer/lyophiliser Alpha 1-2 LD plus (Martin Christ) following manufacturer’s instructions: samples were placed in glass vials with vented rubber caps, placed in a freeze dryer holding tray and placed at −80°C until ready to lyophilise. The freeze dryer machine was cooled down before use and worms were exposed to lyophilisation for 18 hours at −44 °C to −45°C, and pressure at 0.071 to 0.076 mbar. Lyophilized larvae were homogenized in 1 mL PBS using a FastPrep-24 instrument (MP Biomedicals) with 1/4” ceramic sphere in 2 mL tubes for 20 seconds at 6 g.

Five microliters of each extract were mixed with 15 μl of sterile PBS and transferred to a 1.5 mL Eppendorf tube. Micro-Fine U-100 insulin syringes were loaded with the suspension mix and injected intraperitoneally into the midline between the pelvic fins.

### RNA isolation and qPCR experiment

RNA was isolated with Trizol reagent (Invitrogen) according to manufacturer’s instructions. One μg of RNA was reverse-transcribed using M-MLV Reverse Transcriptase Kit (Invitrogen). Real-time PCR was performed using Rox SYBR Green MasterMix dTTP Blue Kit (Takyon) and run on a QuantStudio 6 Flex Real-Time PCR System (Applied Biosystems).

The following primers were used:

*ifng-1-1:* forward, 5’- ACCAGCTGAATTCTAAGCCAA -3’
reverse, 5’-TTTTCGCCTTGACTGAGTGAA -3’
*ifng-1-2:* forward, 5’- CATCGAAGAGCTCAAAGCTTACTA -3’
reverse, 5’-TGCTCACTTTCCTCAAGATTCA -3’
*tnfa:* forward, 5’- TTCACGCTCCATAAGACCCA -3’
reverse, 5’-CAGAGTTGTATCCACCTGTTA -3’
*il13:* forward, 5’- GAAGTGTGAGCATGATTATTTC -3’
reverse, 5’-CTCGTCTTGGTGGTTGTAAG -3’
*il4:* forward, 5’- CCTGACATATATGAGACAGGACACTAC -3’
reverse, 5’-TTACCCTTCAAAGCCATTCC -3’
*il17a/f3:* forward, 5’- AAGATGTTCTGGTGTGAAGAAGTG -3’
reverse, 5’-ACCCAAGCTGTCTTTCTTTGAC -3’
*il22:* forward, 5’- TGCAGAATCACTGTAAACACGA -3’
reverse, 5’-CTCCCCGATTGCTTTGTTAC -3’
*cd3z:* forward, 5’- CCGGTGGAGGAGTCTCATTA -3’
reverse, 5’-CTCCAGATCTGCCCTCCTC -3’

### Alignment and quantification of single-cell RNA-sequencing data

For the samples that were processed using the Smart-seq2 protocol, the reads were aligned to the zebrafish reference genome (Ensemble BioMart version 89) combined with the sequences for EGFP, mCherry, mhc2dab and ERCC spike-ins. Salmon v0.8.2 (Patro et al., 2017) was used for both alignment and quantification of reads with the default paired-end parameters, while library type was set to inward (I) relative orientation (reads face each other) with unstranded (U) protocol (parameter –1 IU).

For the samples that were processed using the Chromium Single Cell 3’ protocol, Cell Ranger v2.1 was used in order to de-multiplex raw base call (BCL) files generated by Illumina sequencers into FASTQ files, perform the alignment, barcode counting, and UMI counting. Ensembl BioMart version 91 was used to generate the reference genome.

### Quality control of single-cell data

For the Smart-seq2 protocol transcript per million (TPM) values reported by Salmon were used for the quality control (QC). Wells with fewer than 900 expressed genes (TPM > 1), or having more than either 60% of ERCC or 45% of mitochondrial content were annotated as poor quality cells. As a result, 322 cells failed QC and 542 single cells were selected for the further study.

Chromium Single Cell 3’ samples were filtered based on the Median Absolute Deviation (MAD) of the distribution of the number of detected genes. In addition, the percentage of mitochondrial content was set to less than 10%. Following QC, 3,211 single cells from the *rag1*^−/−^ PBS-injected sample, 3,626 single cells from the *rag1*^−/−^ A. simplex-injected samples, 3,487 single cells from the *rag1*^−/−^ *V. anguillarum*-injected samples, and 3,756 from the *rag1^+/+^* PBS-injected samples were used in downstream analysis.

### Downstream analysis of Smart-seq2 data

For each of the 542 single cells, counts reported by Salmon were transformed into normalised counts per million (CPM) and used for the further analysis. This was performed by dividing the number of counts for each gene with the total number of counts for each cell and by multiplying the resulting number by a factor of 1,000,000. Genes that were expressed in less than 1% of cells (e.g. 5 single cells with CPM > 1) were filtered out. In the final step we ended up using 16,059 genes across the 542 single cells. The scran R package (version 1.6.7) (L. Lun et al., 2016) was then used to normalise the data and remove differences due to the library size or capture efficiency and sequencing depth.

In order to identify the highly variable genes (HVGs) we utilised the Brennecke Method (Brennecke et al., 2013). We inferred the noise model from the ERCCs and selected genes that vary higher than 20% percentage of variation. This was performed by using the “BrenneckeGetVariableGenes” command of M3Drop v1.4.0 R package setting fdr equal to 0.01 and minimum percentage of variance due to biological factors (minBiolDisp) equal to 0.2. In total, 3,374 were annotated as HVGs.

To verify that all cells were intermixed (in the reconstructed 3D component space) based on their transcriptional similarities and not based on the fish of origin, we used Principal Component Analysis and diffusion maps (destiny R package (version 2.6.1)).

The first 3 diffusion components were clustered using shared nearest neighbour (SNN) modularity optimization-based clustering algorithm implemented by Seurat Package. We used the “FindClusters” command. Three clusters were selected for the further analysis (Supp. Figure 2B).

### BraCeR and TraCeR analysis

We have used TraCeR (Stubbington et al., 2016) and BraCeR (Lindeman et al., 2017) tools in order to reconstruct the sequences of rearranged T and B cell receptor genes (TCR and BCR, respectively), from our Smart-seq2 single-cell RNA-seq data. In order to build combinatorial recombinomes (tracer/bracer build command) for the Danio rerio species, fasta files describing all V, J, C, D sequences were collected from the international ImMunoGeneTics information system (http://www.imgt.org) (Lefranc et al., 2015). For TCR, complete information of the alpha and beta chain location was available, while for the BCR, H and I location sequences were available. Using a threshold of 50 TPMs for gene expression, we identified a total of 244 single cells as TCR positive and 36 as BCR positive.

### Downstream analysis of 10x Genomics data

The downstream analysis of the 10x data was performed using the Seurat (version 2.2.0) and the cellranger (version 1.1.0) R packages. Briefly, raw counts that passed the QC were centered by a factor of 1000 and log transformed. HVGs were detected based on their average expression against their dispersion, by means of the “FindVariableGenes” Seurat command with the following parameters: mean.function equal to ExpMean, dispersion.function equal to LogVMR, x.low.cutoff equal to 0.0125, x.high.cutoff equal to 3, and y.cutoff equal to 0.5. The number of HVGs across samples varied between 1500 and 2500 genes, accordingly.

HVGs were used for the calculation of the Principal Components (PCs) using Seurat’s “RunPCA” command. For the 3D tSNE transformation (“RunTSNE” command) we used PCs with JackStraw statistics lower than 0.01. The later statistics were estimated using the Seurat’s “JackStraw” command with 200 replicate samplings. The proportion of the data that was randomly permuted for each replicate was set to 1%.

Clustering in the 3D tSNE space was performed (pheatmap version 1.0.8) using euclidean distance and centroid linkage. The silhouette scores were used to estimate the optimal number of clusters. However, the final decision on the number of clusters was made on case by case basis. Positive marker genes that expressed in at least half of genes within the cluster were calculated with “FindAllMarkers” Seurat command, using Wilcoxon rank sum test with threshold set to 0.25. DotPlots in Figures 4 and 5 were generated using the Seurat’s “DotPlot” and “SplitDotPlotGG” command, respectively.

### Seurat Alignment Strategy

In order to perform direct comparison of clusters that belong to the same cell type across different conditions, we adopted the Seurat Alignment workflow (Butler et al., 2018). We calculated Highly Variable Genes (HVGs), for each of the three different conditions, and selected 961 HVGs that were expressed in at least 2 datasets. Canonical Correlation Analysis (CCA) was then performed in order to identify shared correlation structures across the different conditions using the “RunMultiCCA” command. Twenty significant CCA components were selected by means of the shared correlation strength, using the “MetageneBicorPlot” command. Aligned CCA space was then generated with the “AlignSubspace” Seurat command. Thirteen Clusters were identified using the shared nearest neighbor (SNN) modularity optimization based clustering algorithm (“FindClusters” command) on the 20 significant CCA aligned components at 0.5 resolution. Dotplot of genes at different clusters on the aligned data was generated by using the SplitDotPlotGG command.

## Acknowledgements

The study was supported by Cancer Research UK grant number C45041/A14953 (to A.C. and E.I.A.), European Research Council project 677501 – ZF_Blood (to A.C. and P.M.S.), EMBO Long-Term Fellowship ALTF-807-2015 (to P.P.H), ANR grant 17-CE15-0017-01 – ZF-ILC (to P.P.H) and ANR-16- CE20-0002-03 (to J.-P.L), H2020-MSCA-IF-2015 grant 708128 – ZF-ILC (to P.P.H) and a core support grant from the Wellcome Trust and MRC to the Wellcome Trust – Medical Research Council Cambridge Stem Cell Institute. The authors would like to thank WTSI Cytometry Core Facility for their help with index cell sorting and the Core Sanger Web Team for hosting the cloud web application. The authors would also like to thank the CRUK Cambridge Institute Genomics Core Facility for their contribution in sequencing the data and WTSI Single Cell Genomics Core Facility for their assistance with 10x experiments.

## Contributions

P.P.H conceived and proposed the original idea of identifying ILCs in zebrafish, under supervision of J.-P. L; C.M.C provided lyophilized *A. simplex*; P.B generated *V. anguillarum* extract. P.P.H and J.-P. L designed, and P.P.H and A.F.R performed inflammation experiments; E.I.A carried out the complete single-cell analysis and created the cloud repository under supervision of A.C; P.M.S performed single-cells experiments, designed and generated all figures with input from A.C.; E.I.A, P.M.S, P.P.H and A.C contributed to the discussion of the results. A.C wrote the manuscript with input from P.M.S, E.I.A, J.-P. L and P.P.H.

## References

Aggad, D., Mazel, M., Boudinot, P., Mogensen, K.E., Hamming, O.J., Hartmann, R., Kotenko, S., Herbomel, P., Lutfalla, G., Levraud, J.-P., 2009. The two groups of zebrafish virus-induced interferons signal via distinct receptors with specific and shared chains. J. Immunol. 183, 3924–31.

Annunziato, F., Romagnani, C., Romagnani, S., 2015. The 3 major types of innate and adaptive cell-mediated effector immunity. J. Allergy Clin. Immunol. 135, 626–35.

Artis, D., Spits, H., 2015. The biology of innate lymphoid cells. Nature 517, 293–301.

Athanasiadis, E.I., Botthof, J.G., Andres, H., Ferreira, L., Lio, P., Cvejic, A., 2017. Single-cell RNA-sequencing uncovers transcriptional states and fate decisions in haematopoiesis. Nat. Commun. 8, 2045.

Björklund, Å.K., Forkel, M., Picelli, S., Konya, V., Theorell, J., Friberg, D., Sandberg, R., Mjösberg, J., 2016. The heterogeneity of human CD127+ innate lymphoid cells revealed by single-cell RNA sequencing. Nat. Immunol. 17, 451–460.

Boehm, T., 2012. Evolution of Vertebrate Immunity. Curr. Biol. 22, R722–R732.

Brennecke, P., Anders, S., Kim, J.K., Kołodziejczyk, A.A., Zhang, X., Proserpio, V., Baying, B., Benes, V., Teichmann, S.A., Marioni, J.C., Heisler, M.G., 2013. Accounting for technical noise in single-cell RNAseq experiments. Nat. Methods 10, 1093–1095.

Brugman, S., 2016. The zebrafish as a model to study intestinal inflammation. Dev. Comp. Immunol. 64, 82–92.

Butler, A., Hoffman, P., Smibert, P., Papalexi, E., Satija, R., 2018. Integrating single-cell transcriptomic data across different conditions, technologies, and species. Nat. Biotechnol. 36, 411–420.

Carmona, S.J., Teichmann, S.A., Ferreira, L., Macaulay, I.C., Stubbington, M.J.T., Cvejic, A., Gfeller, D., 2017. Single-cell transcriptome analysis of fish immune cells provides insight into the evolution of vertebrate immune cell types. Genome Res. 27, 451–461.

Carradice, D., Lieschke, G.J., 2008. Zebrafish in hematology: sushi or science? Blood 111, 3331–3342.

Chang, Y.-J., Kim, H.Y., Albacker, L.A., Baumgarth, N., McKenzie, A.N.J., Smith, D.E., Dekruyff, R.H., Umetsu, D.T., 2011. Innate lymphoid cells mediate influenza-induced airway hyper-reactivity independently of adaptive immunity. Nat. Immunol. 12, 631–8.

Corripio-Miyar, Y., Zou, J., Richmond, H., Secombes, C.J., 2009. Identification of interleukin-22 in gadoids and examination of its expression level in vaccinated fish. Mol. Immunol. 46, 2098–2106.

Dee, C.T., Nagaraju, R.T., Athanasiadis, E.I., Gray, C., Fernandez Del Ama, L., Johnston, S.A., Secombes, C.J., Cvejic, A., Hurlstone, A.F.L., 2016. CD4-Transgenic Zebrafish Reveal Tissue-Resident Th2- and Regulatory T Cell-like Populations and Diverse Mononuclear Phagocytes. J. Immunol. 197, 3520–3530.

Dotiwala, F., Mulik, S., Polidoro, R.B., Ansara, J.A., Burleigh, B.A., Walch, M., Gazzinelli, R.T., Lieberman, J., 2016. Killer lymphocytes use granulysin, perforin and granzymes to kill intracellular parasites. Nat. Med. 22, 210–216.

Eberl, G., Colonna, M., Di Santo, J.P., McKenzie, A.N.J., 2015. Innate lymphoid cells. Innate lymphoid cells: a new paradigm in immunology. Science 348, aaa6566.

Fort, M.M., Cheung, J., Yen, D., Li, J., Zurawski, S.M., Lo, S., Menon, S., Clifford, T., Hunte, B., Lesley, R., Muchamuel, T., Hurst, S.D., Zurawski, G., Leach, M.W., Gorman, D.M., Rennick, D.M., 2001. IL-25 induces IL-4, IL-5, and IL-13 and Th2-associated pathologies in vivo. Immunity 15, 985–95.

Glatzer, T., Killig, M., Meisig, J., Ommert, I., Luetke-Eversloh, M., Babic, M., Paclik, D., Blüthgen, N., Seidl, Seifarth, C., Gröne, J., Lenarz, M., Stölzel, K., Fugmann, D., Porgador, A., Hauser, A., Karlas, A., Romagnani, C., 2013. RORγt+ Innate Lymphoid Cells Acquire a Proinflammatory Program upon Engagement of the Activating Receptor NKp44. Immunity 38, 1223–1235.

Gronke, K., Kofoed-Nielsen, M., Diefenbach, A., 2016. Innate lymphoid cells, precursors and plasticity. Immunol. Lett. 179, 9–18.

Guo, G., Luc, S., Marco, E., Lin, T.-W., Peng, C., Kerenyi, M.A., Beyaz, S., Kim, W., Xu, J., Das, P.P., Neff, T., Zou, K., Yuan, G.-C., Orkin, S.H., 2013. Mapping Cellular Hierarchy by Single-Cell Analysis of the Cell Surface Repertoire. Cell Stem Cell 13, 492–505.

Halim, T.Y.F., Takei, F., 2014. Isolation and Characterization of Mouse Innate Lymphoid Cells, in: Current Protocols in Immunology. John Wiley & Sons, Inc., Hoboken, NJ, USA, p. 3.25.1–3.25.13.

Hernández, P.P., Mahlakõiv, T., Yang, I., Schwierzeck, V., Nguyen, N., Guendel, F., Gronke, K., Ryffel, B., Hölscher, C., Dumoutier, L., Renauld, J.-C., Suerbaum, S., Staeheli, P., Diefenbach, A., 2015. Interferon-λ and interleukin 22 act synergistically for the induction of interferon-stimulated genes and control of rotavirus infection. Nat. Immunol. 16, 698–707.

Hoyler, T., Klose, C.S.N., Souabni, A., Turqueti-Neves, A., Pfeifer, D., Rawlins, E.L., Voehringer, D., Busslinger, M., Diefenbach, A., 2012. The Transcription Factor GATA-3 Controls Cell Fate and Maintenance of Type 2 Innate Lymphoid Cells. Immunity 37, 634–648.

Hurst, S.D., Muchamuel, T., Gorman, D.M., Gilbert, J.M., Clifford, T., Kwan, S., Menon, S., Seymour, B., Jackson, C., Kung, T.T., Brieland, J.K., Zurawski, S.M., Chapman, R.W., Zurawski, G., Coffman, R.L., 2002. New IL-17 family members promote Th1 or Th2 responses in the lung: in vivo function of the novel cytokine IL-25. J. Immunol. 169, 443–53.

Jaitin, D.A., Kenigsberg, E., Keren-Shaul, H., Elefant, N., Paul, F., Zaretsky, I., Mildner, A., Cohen, N., Jung, S., Tanay, A., Amit, I., 2014. Massively parallel single-cell RNA-seq for marker-free decomposition of tissues into cell types. Science 343, 776–9.

Kasheta, M., Painter, C.A., Moore, F.E., Lobbardi, R., Bryll, A., Freiman, E., Stachura, D., Rogers, A.B., Houvras, Y., Langenau, D.M., Ceol, C.J., 2017. Identification and characterization of T reg-like cells in zebrafish. J. Exp. Med. 214, 3519–3530.

Killig, M., Glatzer, T., Romagnani, C., 2014. Recognition strategies of group 3 innate lymphoid cells. Front. Immunol. 5, 142.

L. Lun, A.T., Bach, K., Marioni, J.C., 2016. Pooling across cells to normalize single-cell RNA sequencing data with many zero counts. Genome Biol. 17, 75.

Lane, P.J.L., Gaspal, F.M., McConnell, F.M., Withers, D.R., Anderson, G., 2012. Lymphoid Tissue Inducer Cells: Pivotal Cells in the Evolution of CD4 Immunity and Tolerance? Front. Immunol. 3, 24.

Langenau, D.M., Ferrando, A.A., Traver, D., Kutok, J.L., Hezel, J.-P.D., Kanki, J.P., Zon, L.I., Look, A.T., Trede, N.S., 2004. In vivo tracking of T cell development, ablation, and engraftment in transgenic zebrafish. Proc. Natl. Acad. Sci. U. S. A. 101, 7369–74.

Lazarevic, V., Glimcher, L.H., Lord, G.M., 2013. T-bet: a bridge between innate and adaptive immunity. Nat. Rev. Immunol. 13, 777–789.

Lefranc, M.-P., Giudicelli, V., Duroux, P., Jabado-Michaloud, J., Folch, G., Aouinti, S., Carillon, E., Duvergey, H., Houles, A., Paysan-Lafosse, T., Hadi-Saljoqi, S., Sasorith, S., Lefranc, G., Kossida, S., 2015. IMGT®, the international ImMunoGeneTics information system^®^ 25 years on. Nucleic Acids Res. 43, D413–D422.

Lim, A.I., Li, Y., Lopez-Lastra, S., Stadhouders, R., Paul, F., Casrouge, A., Serafini, N., Puel, A., Bustamante, J., Surace, L., Masse-Ranson, G., David, E., Strick-Marchand, H., Le Bourhis, L., Cocchi, R., Topazio, D., Graziano, P., Muscarella, L.A., Rogge, L., Norel, X., Sallenave, J.-M., Allez, M., Graf, T., Hendriks, R.W., Casanova, J.-L., Amit, I., Yssel, H., Di Santo, J.P., 2017a. Systemic Human ILC Precursors Provide a Substrate for Tissue ILC Differentiation. Cell 168, 1086–1100.e10.

Lim, A.I., Verrier, T., Vosshenrich, C.A., Di Santo, J.P., 2017b. Developmental options and functional plasticity of innate lymphoid cells. Curr. Opin. Immunol. 44, 61–68.

Lindeman, I., Emerton, G., Sollid, L.M., Teichmann, S., Stubbington, M.J.T., 2017. BraCeR: Reconstruction of B-cell receptor sequences and clonality inference from single-cell RNA-sequencing. bioRxiv 185504. https://doi.org/10.1101/185504

Lindemans, C.A., Calafiore, M., Mertelsmann, A.M., O’Connor, M.H., Dudakov, J.A., Jenq, R.R., Velardi, E., Young, L.F., Smith, O.M., Lawrence, G., Ivanov, J.A., Fu, Y.-Y., Takashima, S., Hua, G., Martin, M.L., O’Rourke, K.P., Lo, Y.-H., Mokry, M., Romera-Hernandez, M., Cupedo, T., Dow, L.E., Nieuwenhuis, E.E., Shroyer, N.F., Liu, C., Kolesnick, R., van den Brink, M.R.M., Hanash, A.M., 2015. Interleukin-22 promotes intestinal-stem-cell-mediated epithelial regeneration. Nature 528, 560–564.

Mebius, R.E., Rennert, P., Weissman, I.L., 1997. Developing lymph nodes collect CD4+CD3- LTbeta+ cells that can differentiate to APC, NK cells, and follicular cells but not T or B cells. Immunity 7, 493–504.

Nieuwenhuizen, N., Lopata, A.L., Jeebhay, M.F., Herbert, D.R., Robins, T.G., Brombacher, F., 2006. Exposure to the fish parasite Anisakis causes allergic airway hyperreactivity and dermatitis. J. Allergy Clin. Immunol. 117, 1098–1105.

Patro, R., Duggal, G., Love, M. I., Irizarry, R.A., Kingsford, C., 2017. Salmon provides fast and bias-aware quantification of transcript expression. Nat. Methods 14, 417–419.

Petrie-Hanson, L., Hohn, C., Hanson, L., 2009. Characterization of rag1 mutant zebrafish leukocytes. BMC Immunol. 10, 8.

Picelli, S., Björklund, Å.K., Faridani, O.R., Sagasser, S., Winberg, G., Sandberg, R., 2013. Smart-seq2 for sensitive full-length transcriptome profiling in single cells. Nat. Methods 10, 1096–1098.

Riera Romo, M., Pérez-Martínez, D., Castillo Ferrer, C., 2016. Innate immunity in vertebrates: an overview. Immunology 148, 125–39.

Sanos, S.L., Bui, V.L., Mortha, A., Oberle, K., Heners, C., Johner, C., Diefenbach, A., 2009. RORγt and commensal microflora are required for the differentiation of mucosal interleukin 22–producing NKp46+ cells. Nat. Immunol. 10, 83–91.

Sathe, P., Delconte, R.B., Souza-Fonseca-Guimaraes, F., Seillet, C., Chopin, M., Vandenberg, C.J., Rankin, L.C., Mielke, L.A., Vikstrom, I., Kolesnik, T.B., Nicholson, S.E., Vivier, E., Smyth, M.J., Nutt, S.L., Glaser, S.P., Strasser, A., Belz, G.T., Carotta, S., Huntington, N.D., 2014. Innate immunodeficiency following genetic ablation of Mcl1 in natural killer cells. Nat. Commun. 5, 4539.

Satoh-Takayama, N., Vosshenrich, C.A.J., Lesjean-Pottier, S., Sawa, S., Lochner, M., Rattis, F., Mention, J.-J., Thiam, K., Cerf-Bensussan, N., Mandelboim, O., Eberl, G., Di Santo, J.P., 2008. Microbial Flora Drives Interleukin 22 Production in Intestinal NKp46+ Cells that Provide Innate Mucosal Immune Defense. Immunity 29, 958–970.

Sawa, S., Lochner, M., Satoh-Takayama, N., Dulauroy, S., Bérard, M., Kleinschek, M., Cua, D., Di Santo, J.P., Eberl, G., 2011. RORγt+ innate lymphoid cells regulate intestinal homeostasis by integrating negative signals from the symbiotic microbiota. Nat. Immunol. 12, 320–326.

Scoville, S.D., Mundy-Bosse, B.L., Zhang, M.H., Chen, L., Zhang, X., Keller, K.A., Hughes, T., Chen, L., Cheng, S., Bergin, S.M., Mao, H.C., McClory, S., Yu, J., Carson, W.E., Caligiuri, M.A., Freud, A.G., Freud, A.G., 2016. A Progenitor Cell Expressing Transcription Factor RORγt Generates All Human Innate Lymphoid Cell Subsets. Immunity 44, 1140–50.

Simoni, Y., Fehlings, M., Kløverpris, H.N., McGovern, N., Koo, S.-L., Loh, C.Y., Lim, S., Kurioka, A., Fergusson, J.R., Tang, C.-L., Kam, M.H., Dennis, K., Lim, T.K.H., Fui, A.C.Y., Hoong, C.W., Chan, J.K.Y., Curotto de Lafaille, M., Narayanan, S., Baig, S., Shabeer, M., Toh, S.-A.E.S., Tan, H.K.K., Anicete, R., Tan, E.-H., Takano, A., Klenerman, P., Leslie, A., Tan, D.S.W., Tan, I.B., Ginhoux, F., Newell, E.W., 2017. Human Innate Lymphoid Cell Subsets Possess Tissue-Type Based Heterogeneity in Phenotype and Frequency. Immunity 46, 148–161.

Spits, H., Artis, D., Colonna, M., Diefenbach, A., Di Santo, J.P., Eberl, G., Koyasu, S., Locksley, R.M., J McKenzie, A.N., Mebius, R.E., Powrie, F., Vivier, E., 2013. Innate lymphoid cells — a proposal for uniform nomenclature. Nat. Rev. Immunol. 13.

Stubbington, M.J.T., Lönnberg, T., Proserpio, V., Clare, S., Speak, A.O., Dougan, G., Teichmann, S.A., 2016. T cell fate and clonality inference from single-cell transcriptomes. Nat. Methods 13, 329–332.

Takatori, H., Kanno, Y., Watford, W.T., Tato, C.M., Weiss, G., Ivanov, I.I., Littman, D.R., O’Shea, J.J., 2009. Lymphoid tissue inducer-like cells are an innate source of IL-17 and IL-22. J. Exp. Med. 206, 35–41.

Tang, Q., Iyer, S., Lobbardi, R., Moore, J.C., Chen, H., Lareau, C., Hebert, C., Shaw, M.L., Neftel, C., Suva, M.L., Ceol, C.J., Bernards, A., Aryee, M., Pinello, L., Drummond, I.A., Langenau, D.M., 2017. Dissecting hematopoietic and renal cell heterogeneity in adult zebrafish at single-cell resolution using RNA sequencing. J. Exp. Med. 214, 2875–2887.

Tokunaga, Y., Shirouzu, M., Sugahara, R., Yoshiura, Y., Kiryu, I., Ototake, M., Nagasawa, T., Somamoto, T., Nakao, M., 2017. Comprehensive validation of T- and B-cell deficiency in rag1-null zebrafish: Implication for the robust innate defense mechanisms of teleosts. Sci. Rep. 7, 7536.

Vikström, I.B., Slomp, A., Carrington, E.M., Moesbergen, L.M., Chang, C., Kelly, G.L., Glaser, S.P., Jansen, J.H.M., Leusen, J.H.W., Strasser, A., Huang, D.C.S., Lew, A.M., Peperzak, V., Tarlinton, D.M., 2016. MCL-1 is required throughout B-cell development and its loss sensitizes specific B-cell subsets to inhibition of BCL-2 or BCL-XL. Cell Death Dis. 7, e2345–e2345.

Villani, A.-C., Satija, R., Reynolds, G., Sarkizova, S., Shekhar, K., Fletcher, J., Griesbeck, M., Butler, A., Zheng, S., Lazo, S., Jardine, L., Dixon, D., Stephenson, E., Nilsson, E., Grundberg, I., McDonald, D., Filby, A., Li, W., De Jager, P.L., Rozenblatt-Rosen, O., Lane, A.A., Haniffa, M., Regev, A., Hacohen, N., 2017. Single-cell RNA-seq reveals new types of human blood dendritic cells, monocytes, and progenitors. Science, 356, eaah4573.

Walker, J.A., Barlow, J.L., J McKenzie, A.N., 2013. Innate lymphoid cells — how did we miss them? Nat. Rev. Immunol. 13(2) 75–87.

Wang, L., Bosselut, R., 2009. CD4-CD8 lineage differentiation: Thpok-ing into the nucleus. J. Immunol. 183, 2903–10.

Wang, S., Xia, P., Chen, Y., Yin, Z., Xu, Z., Correspondence, Z.F., Qu, Y., Xiong, Z., Ye, B., Du, Y., Tian, Y., Fan, Z., 2017. Regulatory Innate Lymphoid Cells Control Innate Intestinal Inflammation In Brief Regulatory Innate Lymphoid Cells Control Innate Intestinal Inflammation. Cell 171, 201–216.e18.

Wei, S., Zhou, J., Chen, X., Shah, R.N., Liu, J., Orcutt, T.M., Traver, D., Djeu, J.Y., Litman, G.W., Yoder, J.A., 2007. The zebrafish activating immune receptor Nitr9 signals via Dap12. Immunogenetics 59, 813–821.

Wienholds, E., Schulte-Merker, S., Walderich, B., Plasterk, R.H.A., 2002. Target-Selected Inactivation of the Zebrafish rag1 Gene. Science (80-.). 297, 99–102.

Wilson, N.K., Kent, D.G., Buettner, F., Shehata, M., Macaulay, I.C., Calero-Nieto, F.J., Sánchez Castillo, M., Oedekoven, C.A., Diamanti, E., Schulte, R., Ponting, C.P., Voet, T., Caldas, C., Stingl, J., Green, A.R., Theis, F.J., Göttgens, B., 2015. Combined Single-Cell Functional and Gene Expression Analysis Resolves Heterogeneity within Stem Cell Populations. Cell Stem Cell 16, 712–724.

Wittamer, V., Bertrand, J.Y., Gutschow, P.W., Traver, D., 2011. Characterization of the mononuclear phagocyte system in zebrafish. Blood 117, 7126–7135.

Wolk, K., Kunz, S., Witte, E., Friedrich, M., Asadullah, K., Sabat, R., 2004. IL-22 Increases the Innate Immunity of Tissues. Immunity 21, 241–254.

Yang-Yen, H.-F., 2006 Mcl-1: a highly regulated cell death and survival controller. J Biomed Sci 13(2), 201–4.

Yoder, J.A., Turner, P.M., Wright, P.D., Wittamer, V., Bertrand, J.Y., Traver, D., Litman, G.W., 2010. Developmental and tissue-specific expression of NITRs. Immunogenetics 62, 117–22.

Zheng, G.X.Y., Terry, J.M., Belgrader, P., Ryvkin, P., Bent, Z.W., Wilson, R., Ziraldo, S.B., Wheeler, T.D., McDermott, G.P., Zhu, J., Gregory, M.T., Shuga, J., Montesclaros, L., Underwood, J.G., Masquelier, D.A., Nishimura, S.Y., Schnall-Levin, M., Wyatt, P.W., Hindson, C.M., Bharadwaj, R., Wong, A., Ness, K.D., Beppu, L.W., Deeg, H.J., McFarland, C., Loeb, K.R., Valente, W.J., Ericson, N.G., Stevens, E.A., Radich, J.P., Mikkelsen, T.S., Hindson, B.J., Bielas, J.H., 2017. Massively parallel digital transcriptional profiling of single cells. Nat. Commun. 8, 14049.

Zheng, Y., Valdez, P.A., Danilenko, D.M., Hu, Y., Sa, S.M., Gong, Q., Abbas, A.R., Modrusan, Z., Ghilardi, N., de Sauvage, F.J., Ouyang, W., 2008. Interleukin-22 mediates early host defense against attaching and effacing bacterial pathogens. Nat. Med. 14, 282–289.

